# GWAS of QRS Duration Identifies New Loci Specific to Hispanic/Latino Populations Swenson Hispanic/Latino QRS GWAS

**DOI:** 10.1101/363457

**Authors:** Brenton R. Swenson, Tin Louie, Henry J. Lin, Raú MéndezGiráldez, Jennifer E Below, Cathy C. Laurie, Kathleen F. Kerr, Heather Highland, Timothy A. Thornton, Kelli K. Ryckman, Charles Kooperberg, Elsayed Z. Soliman, Amanda A. Seyerle, Xiuqing Guo, Kent D. Taylor, Jie Yao, Susan R. Heckbert, Dawood Darbar, Lauren E. Petty, Barbara McKnight, Susan Cheng, Natalie A. Bello, Eric A. Whitsel, Craig L. Hanis, Mike A. Nalls, Daniel S. Evans, Jerome I. Rotter, Tamar Sofer, Christy Avery, Nona Sotoodehnia

## Abstract

**Background:** The electrocardiographically quantified QRS duration measures ventricular depolarization and conduction. QRS prolongation has been associated with poor heart failure prognosis and cardiovascular mortality, including sudden death. While previous genome-wide association studies (GWAS) have identified 32 QRS SNPs across 26 loci among European, African, and Asian-descent populations, the genetics of QRS among Hispanics/Latinos has not been previously explored.

**Methods:** We performed a GWAS of QRS duration among Hispanic/Latino ancestry populations (n=15,124) from four studies using 1000 Genomes imputed genotype data (adjusted for age, sex, global ancestry, clinical and study-specific covariates). Study-specific results were combined using fixed-effects, inverse variance-weighted meta-analysis.

**Results:** We identified six loci associated with QRS (*P*<5×10^−8^), including two novel loci: *MYOCD*, a nuclear protein expressed in the heart, and *SYT1*, an integral membrane protein. The top association in the *MYOCD* locus, intronic SNP rs16946539, was found in Hispanics/Latinos with a minor allele frequency (MAF) of 0.04, but is monomorphic in European and African descent populations. The most significant QRS duration association was for intronic SNP rs3922344 (*P*= 8.56×10^−26^) in *SCN5A/SCN10A*. Three additional previously identified loci, *CDKN1A*, *VTI1A*, and *HAND1*, also exceeded the GWAS significance threshold among Hispanics/Latinos. A total of 27 of 32 previously identified QRS duration SNPs were shown to generalize in Hispanics/Latinos.

**Conclusions:** Our QRS duration GWAS, the first in Hispanic/Latino populations, identified two new loci, underscoring the utility of extending large scale genomic studies to currently under-examined populations.

## INTRODUCTION

The duration of the QRS complex, obtained using a resting, standard 12-lead electrocardiogram (ECG), represents the electrical depolarization of the ventricles as an impulse travels through the cardiac conduction system and the ventricular myocardium. Delay in cardiac ventricular conduction results in increased QRS duration, and has been shown to predict heart failure prognosis (1, 2), sudden death (3), and cardiovascular (CV) mortality in patients with and without left ventricular dysfunction, independent of traditional CV risk factors (4). In turn, shortening of the QRS duration with the use of cardiac-resynchronization therapy (CRT) has been shown to decrease heart-failure related events in patients with QRS prolongation (5).

To date, heritability estimates of QRS duration have varied with up to |40% heritability found in more recent studies (6-9). While previous genome-wide association studies have predominantly focused on European populations (10-12), there have been several smaller studies of Asian (13, 14), Pacific Islander (15), and African American populations (16, 17). Collectively, these GWAS analyses have identified 32 SNPs across 26 loci associated with QRS duration. These loci harbor ion channel and transcription factor genes involved in cardiac conduction, including *SCN5A, SCN10A, TBX3, TBX5, TBX20*, and *HAND1* (10-17). To our knowledge, there has been no GWAS performed to study the genetics of QRS duration in Hispanic/Latino ancestry populations. We therefore performed a GWAS of QRS duration in four Hispanic/Latino study populations: the Hispanic Community Health Study/Study of Latinos (HCHS/SOL), the Multi-Ethnic Study of Atherosclerosis (MESA), the Starr County Study (Starr), and the Women’s Health Initiative (WHI).

## RESULTS

Our GWAS included 15,124 Hispanic/Latino individuals from four contributing cohorts. Baseline characteristics varied substantially across the four cohorts. Notably, WHI is a study of women only. The average age of the participants across the four cohorts ranged from 45 to 61 years. The prevalence of hypertension and diabetes across the four cohorts ranged from 26% to 42% for hypertension and 8% to 46% for diabetes (Supplementary Table 1).

### Genome-wide Association Analysis

Following quality control (see Materials and Methods), individual studies contributed between 5.8M and 20.2M individual SNPs, yielding 21.1M combined, unique SNPs overall. Neither the individual studies (λ range=0.96-1.03) nor the combined meta-analysis (λ=1.03) exhibited evidence of test-statistic inflation (Supplementary Figures 1 and 2). SNPs in six loci exceeded the genome-wide threshold for significance (Table 1, Figure 1). Two of the loci (*MYOCD* and *SYT1*) were novel, whereas the remaining four (*SCN5A-SCN10A, HAND1, CDKN1A*, and *VTI1A*) were previously identified in other ethnic groups. There was no evidence of heterogeneity (Cochran’s Q test *P*-value>0.05, Supplementary Table 2), and effect direction was consistent across all contributing studies for all index SNPs (Supplementary Figure 3). All index SNPs were either directly genotyped or were of high imputation quality (Supplementary Table 3).

**Table 1.** Genome-wide significant SNPs identified in a GWAS meta-analysis of n=15,124 participants of Hispanic/Latino ancestry from four studies.

| Locus | Nearest Gene<sup>a</sup> | Index SNP | Chr<sup>b</sup> | A1/A2<sup>c</sup> | Function | CAF<sup>d</sup> | $\beta$ (SE)<sup>e</sup> | P | Multi P<sup>f</sup> |
| --- | --- | --- | --- | --- | --- | --- | --- | --- | --- |
| 1 | SCN5A | rs3922844 | 3 | C/T | Intronic | 0.63 | 1.03 (0.10) | 1.19e-24 | 3.36e-15 |
| 1 | *SCN5A | rs62241190 | 3 | G/A | Intronic | 0.04 | 2.46 (0.27) | 5.82e-20 | 1.09e-12 |
| 1 | *SCN10A | rs10428132 | 3 | T/G | Intronic | 0.38 | 0.79 (0.10) | 1.43e-15 | 3.57e-11 |
| 1 | *SCN5A | rs9856387 | 3 | C/T | Intronic | 0.73 | 0.76 (0.11) | 2.12e-12 | 1.68e-08 |
| 2 | HAND1 | rs13165478 | 5 | G/A | Intergenic | 0.67 | 0.68 (0.10) | 2.69e-11 | -- |
| 3 | CDKN1A | rs3176326 | 6 | A/G | Intronic | 0.17 | 1.15 (0.13) | 1.54e-19 | 4.26e-13 |
| 3 | **SPRK1 | rs2395642 | 6 | T/C | Intronic | 0.16 | 0.93 (0.13) | 4.71e-09 | 3.94e-06 |
| 4 | VTI1A | rs7906312 | 10 | A/C | Intronic | 0.19 | 0.77 (0.13) | 8.14e-10 | -- |
| 5 | SYT1 | rs4842438 | 12 | C/A | Intronic | 0.93 | 1.04 (0.19) | 4.24e-08 | -- |
| 6 | MYOCD | rs16946539 | 17 | T/C | Intronic | 0.06 | 1.28 (0.22) | 1.74e-09 | -- |

<sup>a</sup>\* Denotes a secondary signal. \*\* Denotes a secondary SNP no longer genome-wide significant after multi-SNP testing at the locus.
<sup>b</sup>Chr: Chromosome
<sup>c</sup>A1/A2: Coded/non-coded alleles
<sup>d</sup>CAF: Coded allele frequency
<sup>e</sup> $\beta$ : Effect estimate measured in milliseconds.
<sup>f</sup>P for the association of index SNP with QRS duration, upon fixed effects meta-analysis using a model which includes all index SNPs in the same locus.

**Figure 1.**
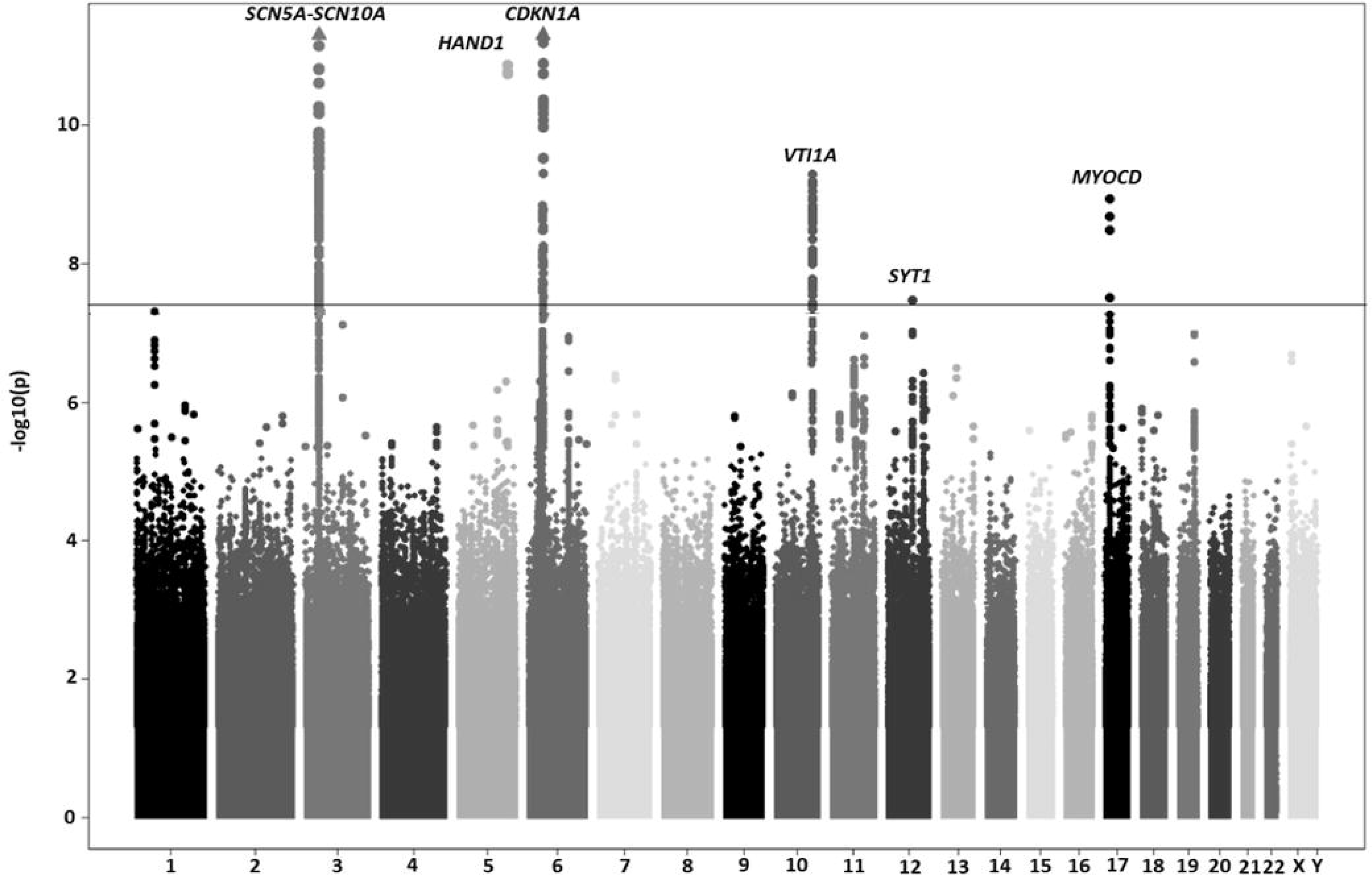
Manhattan plot of SNP-QRS associations. Manhattan plot showing the association of SNPs with QRS duration in the GWAS meta-analysis containing 15,124 individuals of Hispanic/Latino ethnicity. The horizontal line represents the genome-wide significance threshold of (*P*=5E-08). SNPs mapping to 6 loci exceeded the GWAS threshold for significance.

### Novel Associations

The meta-analysis identified two novel loci (*MYOCD* and *SYT1)* associated with QRS duration. The index SNP (rs16946539) in *MYOCD* (myocardin, a nuclear protein found in cardiac and smooth muscle), as well as the only SNP (rs139859815) in high LD (r^2^>0.5) with it, are monomorphic in the European-descent and African-descent 1000 Genomes super populations (Supplementary Table 4).

The second novel locus was on chromosome 12 near *SYT1* (synaptotagmin-1), an integral membrane protein of synaptic vesicles that responds to calcium signaling. The index SNP in *SYT1* (rs4842438) was examined in both European and African ancestry GWAS efforts, but failed to reach nominal significance (*P*>0.05; Figure 3). Indeed, the effect size of rs4842438 in Hispanics/Latinos (beta=1.04 ms) is statistically larger than in European-descent (beta=0.13 ms) and African-descent (beta=0.00 ms) individuals (*P* for difference=5.69×10^−5^ and 8.65×10^−6^, respectively, Figure 3, Supplementary Tables 5, 6, and 7). Moreover, the broad LD pattern seen in Hispanics/Latinos is entirely absent in Europeans and African Americans (Supplementary Figure 4E). The lack of associations among European and African descent individuals is not explained by lack of power due to smaller sample size or lower MAF (European MAF=0.06; African MAF=0.20; Hispanic/Latino MAF=0.07).

### SCN5A-SCN10A

The most significant association with QRS duration was found in chromosome region 3p22 (rs3922844) in locus 1, bridging *SCN5A* and *SCN10A*, two adjacent cardiac sodium-channel genes (Figure 2). This SNP had previously been found to be genome-wide significant among European and African American descent individuals (Supplementary Tables 5 and 6). Similar to our findings in Hispanics/Latinos, rs3922844 is the most significantly associated QRS SNP in African Americans (Figure 2). In contrast, among European-descent individuals, the strongest SNP association (rs1601957) resides in an intron of the *SCN10A* gene. The effect size of rs3922844 in Hispanics/Latinos is statistically larger than in Europeans (1.32 ms vs 0.56 ms decrease in QRS duration, respectively, *P* for difference=5.8×10^−5^), and is nearer to the effect size seen among African Americans (0.94 ms, Figure 3 and Supplementary Tables 5, 6, and 7). Conditional analyses at the *SCN5A-SCN10A* locus among Hispanics/Latinos revealed three additional independent genome-wide significant secondary signals (Table 1). Two of the secondary index SNPs (rs62241190 and rs9856387) were not tested in either the European or African American GWAS, as those efforts were based on HapMap rather than 1000 Genomes imputation. There are no SNPs in these two populations that are in high LD (r^2^>0.75) with rs62241190 and rs9856387. Therefore, whether these SNPs are also significantly associated with QRS duration among nonHispanic/Latino groups is unknown.

**Figure 2.**
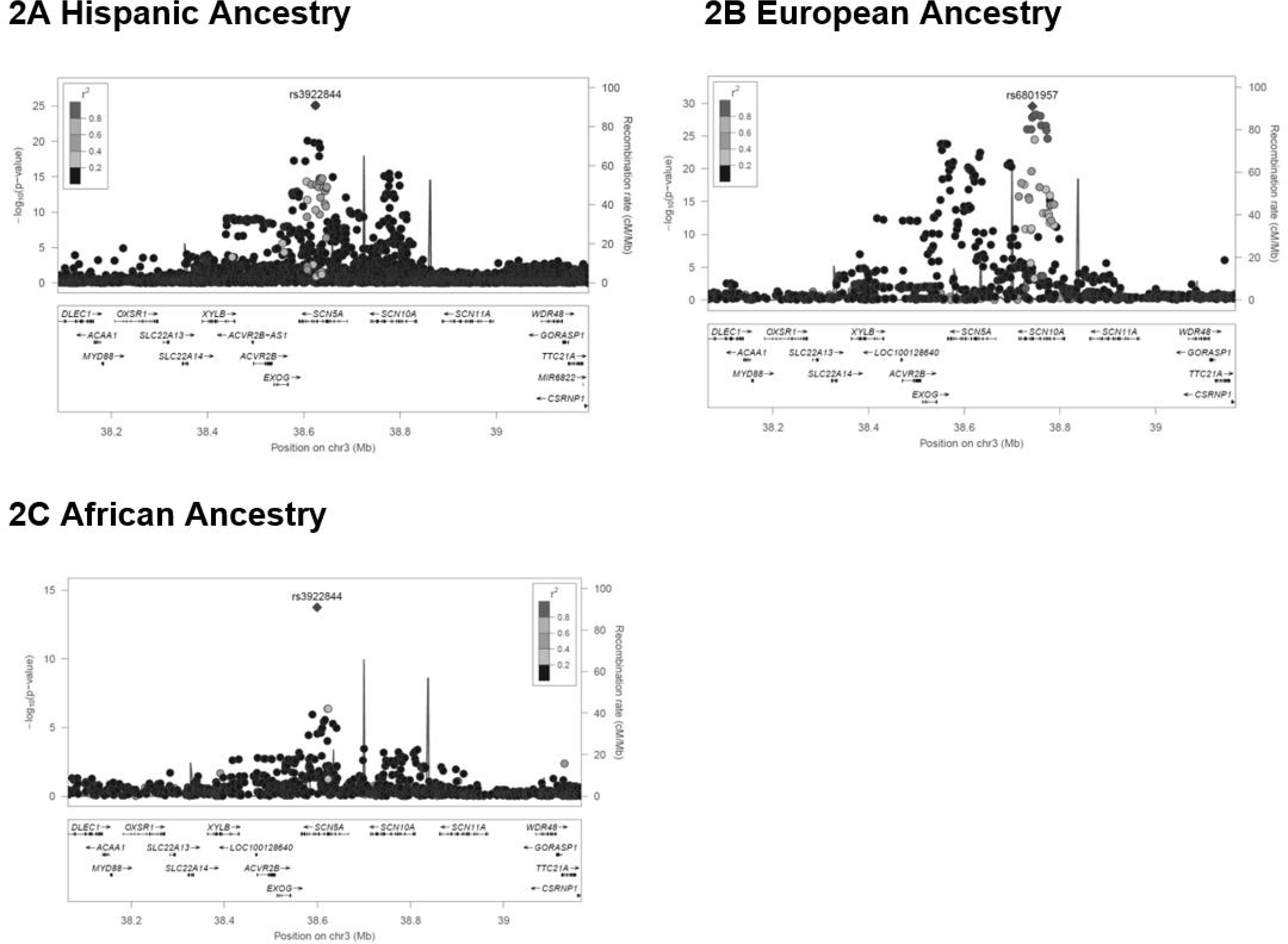
LocusZoom plots of *SCN5A-SCN10A*. A (top-left) – Hispanic/Latino GWAS. B (top right) – European GWAS. C (bottom left) – African American GWAS. The most significant SNP identified in the Hispanic Latino and African American GWAS was rs3922844 in *SCN5A*. The most significant SNP in the European GWAS was rs6801957 in *SCN10A*.

**Figure 3.**
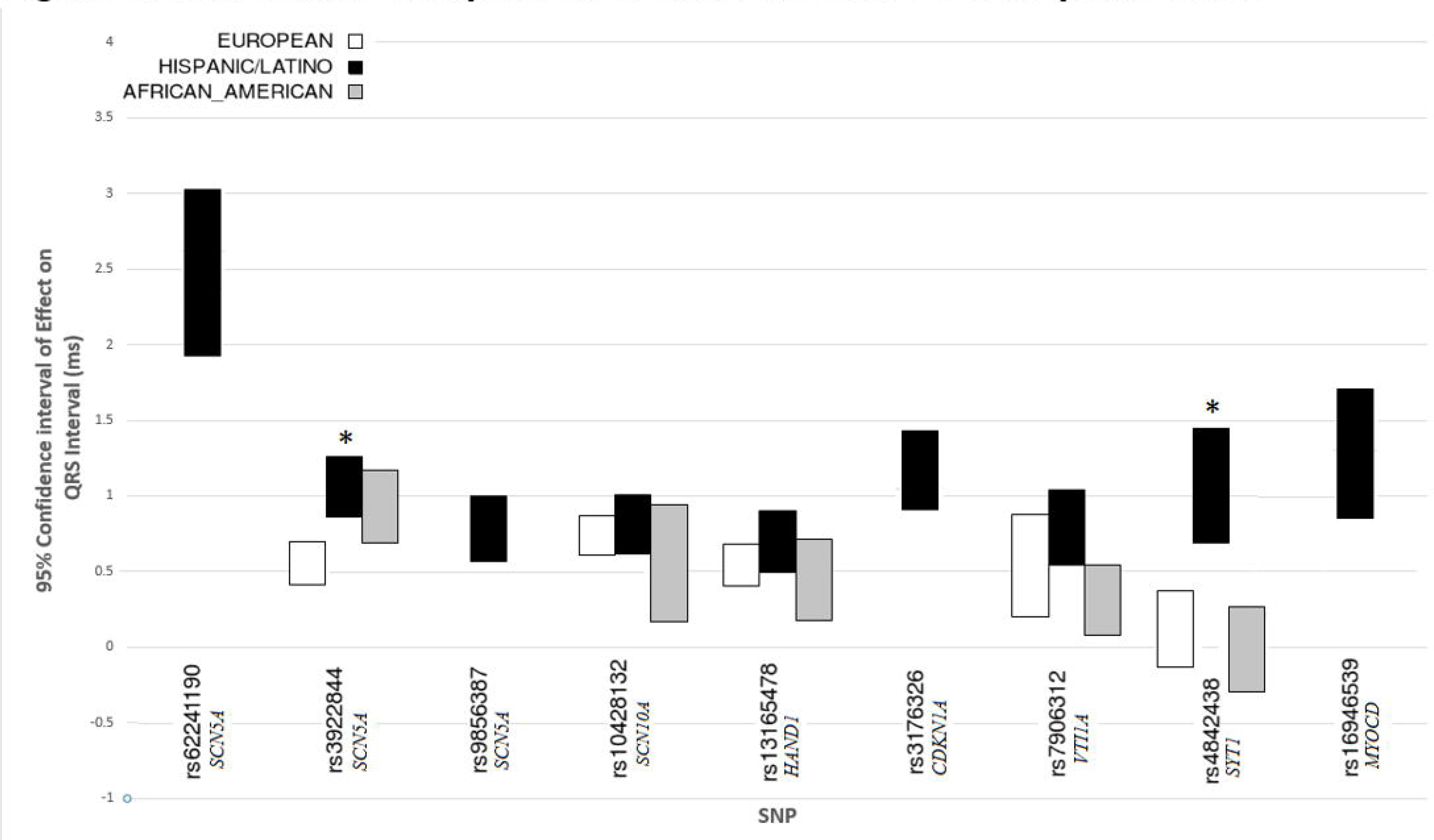
Transethnic comparison of QRS effect sizes for Hispanic/Latino SNPs. Comparison of effect sizes and 95% confidence interval for the 9 index SNPs that were genome-wide significant in the Hispanic/Latino QRS duration GWAS, and the corresponding effect sizes for those SNPs in the European and African American GWAS. Note that rs10428132 was not directly measured in the European or African American studies, but instead a SNP in perfect LD (r^2^=1) was used (rs6800541). Other SNPs that were not directly measured in Europeans or African Americans are not presented (rs62241190, rs9856387, rs3176326, and rs16946539). * Refers to SNPs where the difference in effect size between two ethnic groups was significant at the Bonferroni corrected *P*-value. Two SNPs showed larger effects in Hispanics/Latinos than in European-descent individuals: rs3922844 in *SCN5A* and rs4842438 in *SYT1*. See Supplementary Table 7 for additional details.

### Additional Hispanic/Latino Associations in Known Loci

Three additional previously discovered SNP-QRS associations were also found in Hispanics/Latinos. These include intronic SNPs within *CDKN1A* and *VTI1A*, and an intergenic SNP near *HAND1*. While the Hispanic/Latino index SNP in *CDKN1A* (rs3176326) was not directly evaluated in the European or African American GWAS, it was in high LD with a SNP that had been found to be highly significant in the previous European ancestry GWAS (r^2^=0.71 with rs9462210). Interestingly, a conditional analysis of the *CDKN1A* locus revealed a suggestive secondary signal located in an intron of *SPRK1* approximately 1 Mb upstream from the primary *CDKN1A* signal (rs2395642, *P*=4.58×10^−6^ in conditional analyses; Table 1 and Supplementary Figure 4C). The index SNP in *VTI1A* (rs7906312) is a novel SNP within a known locus, but it is not in high LD with the previously known SNP associated with QRS duration in Europeans (Supplementary Table 5 and Supplementary Figure 4D). Findings for *HAND1* (rs13165478) show that five SNPs in the region, all in very high LD with each other, were significantly associated with QRS duration in this 1000 Genomes imputation analysis, with no other SNPs near significance. This same haplotype was also identified as significant in those of European descent (Supplementary Figure 4B).

### Transethnic Analyses: Generalization

We examined association of 32 index SNPs from published GWAS analyses (27 European,1 African-American, 2 East Asian, and 2 from a meta-analysis of the European and African American GWAS results) for association with QRS duration among Hispanics/Latinos (11, 13, 17) (Supplementary Table 8, Figure 4). Of the 32 previously identified independent SNPs, 27 generalized in Hispanics/Latinos (r-value <0.05). These included 26 of the 27 SNPs from the European GWAS, with rs1362212 as the exception (Supplementary Table 8).

**Figure 4.**
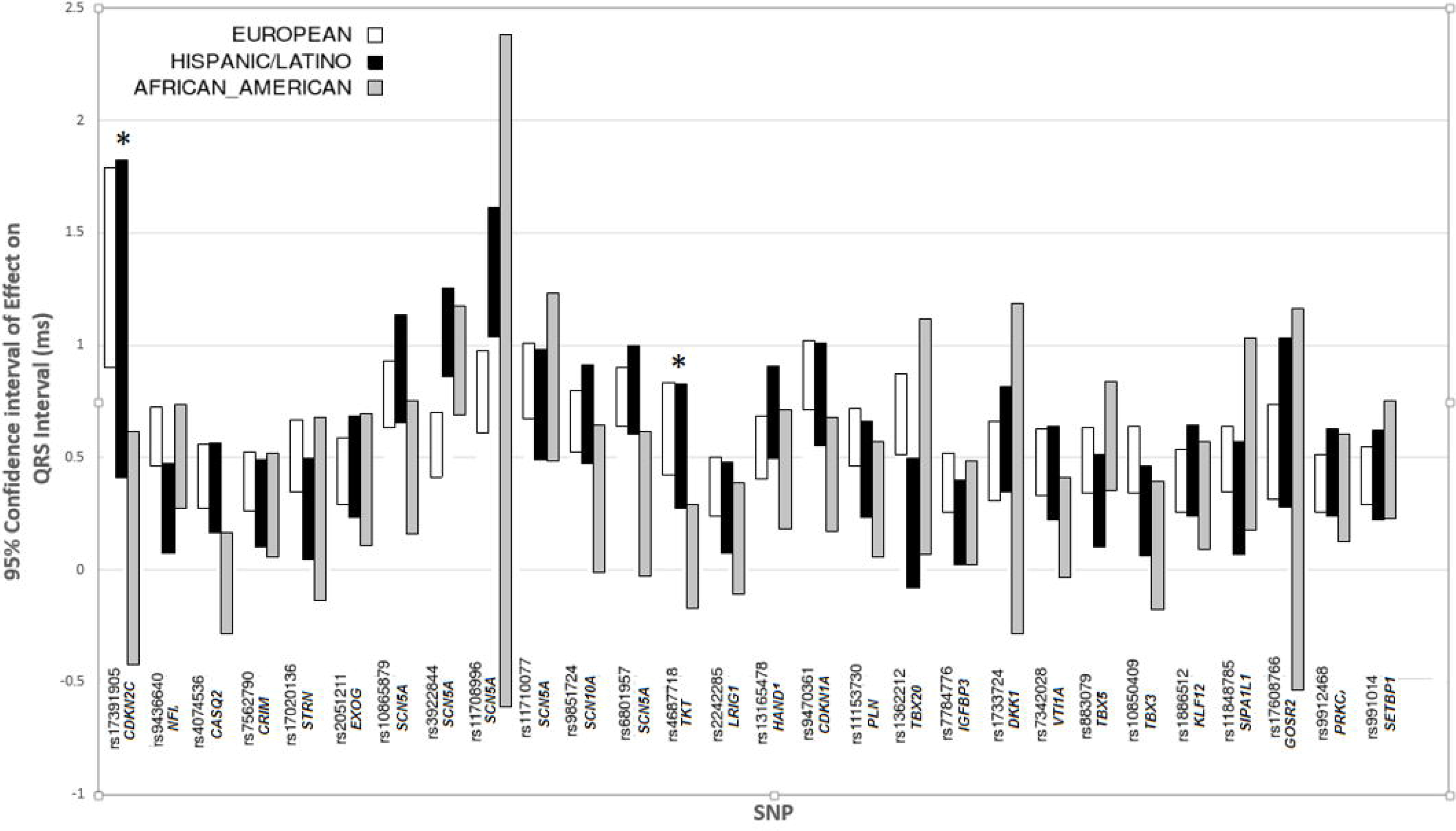
Transethnic comparison of QRS effect sizes for previously known SNPs. Comparison of effect sizes and 95% confidence intervals for 28 previously discovered SNPs for QRS duration across European, Hispanic/Latino, and African American GWAS results. Largely, findings are similar across ethnic groups. * Refers to SNPs where the difference in effect size between two ethnic groups was significant at the Bonferroni corrected *P*-value. See Supplementary Table 7 for additional details.

### Transethnic Analyses: Effect Size

We then compared the effect sizes of the independent index SNPs identified in Hispanic/Latino, European descent, and African descent populations (using the current analysis for Hispanic/Latinos, and published meta-analyses for European and African descent individuals; Figures 3 and 4). Of note, due to differences in imputation sets (HapMap imputation in those of European and African descent versus 1000 genomes imputation in the current analysis), not all Hispanic/Latino SNPs were available in European and African-descent individuals. Hence, there are missing European and African-descent data for 4 SNPs because 3 had no proxy in high LD (r^2^>0.9) with the index Hispanic/Latino SNP, and one was monomorphic in these other populations. After Bonferroni correction, 4 of the 29 independent SNPs showed evidence of significant differences in genotype-phenotype effect sizes (Figures 3 and 4, Supplementary Table 7).

### Cross Phenotype Analyses

The 9 index SNPs identified in the Hispanic/Latino QRS duration GWAS were also examined for their association with other ECG phenotypes: QT duration (18), PR duration (19), heart rate (20), and a measure of heart rate variability, the standard deviation of normal to normal R-R intervals (SDNN) (20) (Table 2, Supplementary Table 9). The 4 SNPs in the *SCN5A-SCN10A* locus were genome-wide significantly associated with both QT and PR interval duration, with SNPs that prolong QRS also prolonging the PR interval but conversely shortening QT duration. This pattern was also observed in the *CDKN1A* locus. However, the index SNPs rose only to a Bonferroni corrected significance level for QT duration, and fell just under significance for PR duration (*P*=1.44E-03). Intriguingly, the novel *MYOCD* locus was significantly associated with PR duration at the Bonferroni corrected significance level, with the SNP that prolongs QRS duration shortening PR duration. None of the QRS index SNPs were significantly associated with heart rate or SDNN.

**Table 2.** Pleiotropic Analyses. Comparison of the effect size and significance level of QRS prolonging index SNPs with QT and PR duration in Hispanics/Latinos. There was no association of QRS SNPs with heart rate or heart rate variability (SDNN). Only significant results (*P*<0.05) are shown.

| Nearest Gene | SNP | QRS $\beta$ (ms) | QRS $P$ | PR $\beta$ (ms) | PR $P^a$ | QT $\beta$ (ms) | QT $P^a$ |
| --- | --- | --- | --- | --- | --- | --- | --- |
| SCN5A | rs62241190 | **2.46** | 5.82E-20 | **4.72** | 2.90E-12 | **-3.58** | 1.83E-09 |
| SCN5A | rs3922844 | **1.03** | 1.19E-24 | **3.39** | 3.57E-11 | **-1.77** | 9.52E-16 |
| SCN5A | rs9856387 | **0.76** | 2.12E-12 | **1.88** | 8.90E-12 | **-1.42** | 3.43E-09 |
| SCN10A | rs10428132 | **0.79** | 1.43E-15 | **3.81** | 4.89E-53 | **-1.29** | 3.04E-09 |
| HAND1 | rs13165478 | **0.68** | 2.69E-11 | - |  | - |  |
| CDKN1A | rs3176326 | **1.15** | 1.54E-19 | **1.02** | 1.44E-03 | **-1.1** | 8.59E-05 |
| VTI1A | rs7906312 | **0.77** | 8.14E-10 | - |  | - |  |
| SYT1 | rs4842438 | **1.04** | 4.24E-08 | - |  | - |  |
| MYOCD | rs16946539 | **1.28** | 1.74E-09 | **-2.28** | 1.82E-05 | - |  |

<sup>a</sup>Bonferroni corrected significance: $P < 1.39E-03$ for 36 tests

### Functional Annotation

The function for all 9 index SNPs was investigated using the HaploReg 4.1 web server (21). Functional information was obtained for 3 loci: *SCN5A-SCN10A, CDKN1A*, and *MYOCD*. SNPs in *CDKN1A* showed evidence for activating transcription in heart tissues, including fetal heart tissue, the right atrium, the right ventricle, and the left ventricle. *SCN5A-SCN10A* and *MYOCD* SNPs were identified as possible enhancers of transcription in the same heart tissues (Supplementary Table 10).

## DISCUSSION

Our GWAS meta-analysis of four cohorts containing 15,124 individuals of Hispanic/Latino ethnicity found 9 index SNPs across 6 loci with genome-wide significant associations with QRS duration. Two loci were novel (*MYOCD* and *SYT1)* and four of the loci (*SCN5A-SCN10A, CDKN1A, HAND1*, and *VTI1A*) were previously identified in QRS GWAS analyses of European- (11) and African-descent (17) individuals. This is the first GWAS of QRS duration in Hispanics/Latinos, a genetically admixed group, comprised of European, African, and Native American ancestry populations, for whom genetic as well as non-genetic determinants of cardiovascular risk remain under-examined (22, 23).

We identified two novel loci associated with QRS duration among those of Hispanic descent: *MYOCD* and *SYT1*. *MYOCD* codes for a nuclear protein (myocardin) expressed in cardiomyocytes and smooth muscle cell-containing tissues. *MYOCD* has been shown to be essential for maintaining adult heart function (24). Mice in which *MYOCD* is postnatally knocked down develop dilated cardiomyopathy and fatal heart failure (25). The index SNP in *MYOCD* (rs16946539) is more common among Hispanics/Latinos (defined by the Ad Mixed American 1000 Genomes population, MAF=0.04) than any of the other 1000 Genomes super populations. Moreover, the *MYOCD* SNP is monomorphic among European- and African-descent individuals (European MAF=0.0; Asian MAF=0.02; African MAF=0.0). Therefore, a QRS GWAS in Hispanics/Latinos is uniquely advantageous in uncovering this genotype-phenotype association. This finding illustrates one of the main imperatives for conducting genetic studies in diverse and under-examined populations, both for ECG traits, and for other traits. Because underlying genetic architecture can differ across racial and ethnic populations, examining new populations may uncover novel genotype-phenotype associations.

The second novel locus found in this study involved an intronic SNP in *SYT1*. Interestingly, a different synaptotagmin gene, *SYT10*, was previously associated with heart rate (26). The *SYT1* intronic SNP was not associated with heart rate in our study. It is noteworthy that this SNP in *SYT1* showed no evidence of association with QRS among European and African-descent individuals. Further studies are warranted to validate this novel association in additional Hispanics/Latinos populations.

The most significant association signal was found in and surrounding two voltage-gated sodium channel genes: *SCN5A* and *SCN10A*. Whereas *SCN5A* is the canonical cardiac sodium channel responsible for cellular depolarization and enables conduction of the electrical signal, *SCN10A* appears to be particularly enriched in the specialized *His Purkinje* conduction fibers (11). Transethnic analyses intriguingly show that the regional association plot for QRS in Hispanics/Latinos more closely resembles that of African Americans than plots of European-descent individuals, with rs3922844 being the most significant SNP in the region among Hispanics/Latinos and African Americans.

The generalization study of the 28 independent index SNPs from QRS GWAS analyses in Hispanic/Latino, European, and African ancestry individuals shows remarkable consistency in the magnitude and direction of the effects overall. While, the genetic architecture of QRS duration in the three ethnic groups is largely comparable, differences are also present, such as the larger effect size at rs3922844 among Hispanics/Latinos than European descent individuals.

Several limitations deserve consideration. First, our sample size, while representing the largest GWAS performed in Hispanics/Latinos, was nonetheless small. Additional Hispanic/Latino cohort studies are needed to extend these findings. Nonetheless, despite its relatively small sample size, novel associations were identified, and known associations were confirmed in a new ethnic population. Larger sample size may reveal more genome-wide significant loci. For example, a SNP in the previously identified *NFIA locus* fell just short of genome-wide significance in Hispanics/Latinos (*P*=5.1×10^−8^). Furthermore, while we excluded individuals with QRS duration longer than 120 ms in order to exclude individuals who have conduction defects and/or bundle branch blocks due to acquired heart disease, interesting genetic associations may be missed by this approach.

In conclusion, our findings indicate that the genetics of QRS duration are largely similar between ethnic groups. However, important differences do exist, illustrated, for instance, by the novel genome-wide significant SNP in *MYOCD* that is monomorphic in both Europeans and African Americans. Our study underscores the importance of conducting genetic studies in diverse and under-examined populations, such as Hispanics/Latinos, to uncover novel loci.

## MATERIALS AND METHODS

### Study Population – Primary meta-analysis of individuals with Hispanic/Latino ancestry

Our meta-analysis included 15,124 participants of self-identified Hispanic/Latino descent from the following four studies: the HCHS/SOL (n=11,566), the Multi-Ethnic Study of Atherosclerosis (MESA, n=1431), the Starr County Study (n=582), and the WHI (n=1545) (see Supplementary Material for cohort description and Supplementary Table 1 for baseline characteristics by cohort). Ancestry was confirmed through principal components analysis, and a small number of genetic outliers (individuals determined to be of primarily Asian ancestry) were excluded.

### Study Population – Comparison meta-analyses of individuals with European and African ancestry

Comparisons of results were made between the Hispanic/Latino meta-analysis, and two published meta-analyses of QRS duration in individuals of European (n=40,407) and African American (n=13,301) ancestries. Details of these studies can be obtained from their original publications (11, 17).

### Electrocardiography

Participants in each of the four cohorts underwent a standard 12-lead ECG by a certified technician (see Supplementary Table 11). Participants were excluded from further analysis if they had any of the following: poor quality ECGs, atrial flutter or fibrillation, a ventricular paced rhythm, QRS duration ≥120 ms, Wolff-Parkinson-White on ECG, a history of previous myocardial infarction or heart failure, or were taking class I or class III antiarrhythmic medications.

### Genotyping and imputation

HCHS/SOL participants were genotyped on an Illumina custom array that included the Illumina Omni 2.5M array (HumanOmmni2.5-8v1-1) and an additional |150,000 SNPs. The additional SNPs were chosen to contain markers relevant to Hispanic/Latino ancestry, markers informative of Native American ethnicity, and significant loci from previous association studies (21). MESA, WHI, and Starr County participants were genotyped using the Affymetrix Genome-Wide Human SNP Array 6.0. SNP genotyping inclusion criteria varied slightly across studies. After individual cohort genotype QC, imputation based on the 1000 Genomes phase 1 reference panel (27) was performed resulting in roughly 38 million SNPs (Supplementary Table 11).

### Statistical Analysis

To assess the association between genotype and QRS duration, individual cohort studies used additive genetic linear regression models, either in a regression model (MESA, WHI, Starr County) or a mixed model (HCHS/SOL, to account for relatedness and shared environment between individuals). The two methods estimate the same effect. Models were adjusted for age, sex, heart rate, systolic blood pressure, BMI, height, study site/region, and principal components of genetic ancestry. After we received the results from the studies, we applied an individual cohort QC filter, which excluded SNPs with low imputation quality (<0.30) or small effective sample sizes for each individual SNP (*effN* <30), with *effN* = 2 × *MAF* × (*MAF* –1) × *N* × *Imputation Quality;* where N is the number of participants. Each cohort contained between |6M and |20M imputed SNPs, after applying this filter.

Results were combined using fixed-effects inverse variance meta-analysis using the METAL software package (28), using genomic control for summary statistics to reduce test-statistic inflation. Study heterogeneity was evaluated using the Cochran Q test. Approximately 21M unique SNPs were contained in the meta-analysis. Results were considered genome-wide significant for *P*-values < 5×10^−8^. Secondary signals were identified using iterative rounds of conditional analysis, with adjustment for additional Hispanic/Latino index SNPs in the model, until there were no SNPs found to be genome-wide significant.

### Transethnic Analyses: Generalization

Previous meta-analyses in European, East Asian, African American ancestry populations identified a total of 32 genome-wide significant SNPs. To assess whether these significant findings generalize to Hispanics/Latinos, we used the method of Sofer *et al* (29). An association is considered generalized if a significant effect in the same direction exists in both the non-Hispanic/Latino discovery population as well as the Hispanic/Latino population. This method controls the false discovery rate (FDR) of the generalization null hypotheses, and generates an r-value for each SNP (with r-values < 0.05 showing evidence that an association is generalizable under FDR control at the α=0.05 level.)

### Transethnic Analyses: Effect Size

Comparisons of effect size differences of SNPs on QRS duration across the Hispanic/Latino, European, and African American GWAS were done using a procedure analogous to Welch’s *t*-tests for each of the 33 independent SNPs that were identified as having a primary or secondary independent association in any of those studies. For the purposes of Bonferroni correction, there were 29 independent SNPs available for testing in all 3 cohorts. Some SNPs were not available in all cohorts due to use of different imputation panels. Comparisons for 29 SNPs among 3 studies resulted in 87 tests. Therefore, differences in effect size were determined to be significant when *P*<5.75E-04.

### Cross Phenotype Analysis

Index SNPs discovered in the QRS duration analysis were also examined for associations with other ECG phenotypes, including QT, PR, heart rate, and SDNN. These GWAS efforts were based upon the same underlying Hispanic/Latino cohorts as the QRS duration GWAS. However, due to different inclusion/exclusion criteria, differences in the study samples do exist between studies. Full details of these studies can be obtained from their original publications (18-20). Significance for these other traits exceeded either: a genome-wide significance threshold (*P<5.0E-08)*; Bonferroni corrected significance (*P<*1.39E-03 for 36 tests); or nominal significance (*P<0.05)*.

### Functional Annotation

The HaploReg v4.1 online web resource was used to functionally annotate genome-wide significant SNPs (21). HaploReg utilizes data obtained from the ENCODE (30) and RoadMap projects (31), to give information on how SNPs might alter gene expression in diverse tissue types. We restricted our analysis to only specific heart tissues – namely, fetal heart tissue, the right atrium, the right ventricle, and the left ventricle. Based on the chromatin-15 state model, we summarized the potential function of SNPs in the genome-wide significant loci in each of the different heart tissues. For each loci, we examined primary SNPs, secondary signal SNPs, and all other SNPs in high LD (r^2^>0.80) with these SNPs. The LD structure pattern used for this analysis was the 1000 Genomes AMR Phase-1 super-population.

## ACKNOWLEDGMENTS

We acknowledge the work of the CHARGE QRS GWAS Consortium and the CARe-COGENT African-American QRS Consortium for their work on the respective European and African American GWAS studies referenced here. Full membership for these two groups is provided in the Supplemental materials.

*Hispanic Community Health Study/Study of Latinos (HCHS/SOL):* We thank the participants and staff of the HCHS/SOL study for their contributions to this study.

*Multi-Ethnic Study of Atherosclerosis (MESA):* We also thank the other investigators, the staff, and the participants of MESA for their valuable contributions. A full list of participating MESA investigators and institutions can be found at http://www.mesanhlbi.org.

Starr County: We thank the field staff in Starr County for collection of these data and are grateful to the study participants who gave their time and contributed to the study. These studies (protocol SPH-02-042) were approved by the University of Texas Health Science Center at Houston’s Committee for the Protection of Human Subjects and carried out in a manner consistent with the Declaration of Helsinki. The studies were explained to all participants and written informed consent obtained. Genotyping services were provided by the Center for Inherited Disease Research (CIDR).

Women’s Health Initiative Clinical Trial (WHI CT): We thank the study participants and staff for contributions to the study.

